# A trait-based framework for quantifying arthropod invasion potential: Predictive modeling with *Tropilaelaps* mites as a case study

**DOI:** 10.64898/2026.05.06.723306

**Authors:** Carmen Black, Treson Thompson, Madison Sankovitz, Samuel Ramsey

## Abstract

Over the past decade, the global rise in invasive species has accelerated at an unprecedented rate, intensifying threats to ecosystems, human health, and economies worldwide. Newly invasive taxa, such as *Tropilaelaps* mites, are of particular concern for apiculture and agroecosystems. Despite growing concern about the spread of *Tropilaelaps* mites and other arthropods, limited resources are available to assess their invasive potential. We characterized 118 invasive arthropod species using available literature to identify key biological and ecological traits associated with invasive potential. We developed predictive generalized linear mixed models (GLMMs) to determine the traits most important for predicting invasive potential (number of invaded regions), and the top-performing models were subsequently applied to *Tropilaelaps mercedesae*. Several traits were identified as significant predictors of invasiveness, including the degree of human association, resilience at small population sizes, diet breadth, maximum annual number of generations, altitude range, and the interaction between human association and temperature range. Notably, *T. mercedesae* was predicted to be capable of invading 160 regions, ranking it within the top 10% most invasive species among those evaluated (12th out of 119), ranked just below the cosmopolitan *Varroa destructor* mite. These findings position *T. mercedesae* as a high-risk, yet under-recognized, invasive threat. Collectively, this demonstrates the power of predictive trait-based modeling to inform invasion risk prior to widespread establishment and underscores the urgency of reallocating resources toward surveillance, research, and proactive management strategies rather than relying on costly, often ineffective post-establishment eradication.

## Introduction

Biological invasions by non-native species present a major and growing challenge for ecosystems, economies, and human societies worldwide. In the United States alone, the cumulative economic and environmental costs of invasive species from 1960 to 2020 have been estimated at $4.52 trillion, with annual losses ranging from approximately $20 to $120 billion (Pimentel et al. 2005; Fantle-Lepczyk et al. 2022). A recent synthesis of invasive species impacts in Australia found that, across losses and management activities, invasives have cost the country 389.6 billion AUD (or 298.6 billion USD in 2017 dollars) (Bradshaw et al. 2021; USDA 2025). However, invasive species can have far-reaching consequences for invaded environments, including ecosystem degradation and fragmentation (Hoffmeister et al. 2005), biodiversity loss (Linders et al 2019), threats to native species (Bennett et al. 2011), risks to human health (Neil & Arim 2011),substantial economic impacts (Fantle-Lepczyk et al. 2022), and can have many cross-ecosystem effects (Peller & Altermatt 2024). For example, the invasion of the forest pest emerald ash borer in North America has led to the destruction of millions of ash trees-permanently altering ecosystems, increasing wildfire risk, and leading to severe economic loss (Jones & McDermott 2018). As well, the invasion of Vespula wasps in North America has posed threats to human health - due to their intense stings-and agriculture-due to their predation of honey bees (Scarparo et al. 2021; Sankovitz et al. 2023; Taylor et al. 2024). Hence, management of invasive species matters not only for biodiversity and ecosystem integrity, but also for human well-being, agriculture, and economic stability. As such, developing effective, targeted exclusion plans before the introduction of invasive organisms is imperative. However, designing these plans for every potential invasion is impractical. Our planet, once composed of discrete ecosystems, has been increasingly globalized, providing innumerable opportunities for invasion. Here, we implement a predictive model to target species that have high trait-based invasion potential. As a case study, we use data on invasive arthropod species to predict the invasion potential of *Tropilaelaps* mites (Class: Arachnida, Order: Mesostigmata), an emerging threat to economies supported by the western honey bee (*Apis mellifera*).

Invasion biology as a field has many conceptually overlapping and ill-defined terms.

Here, we attempt to clearly define some of these terms to avoid confusion. A native species is one that naturally occurs within its historic or indigenous range, typically defined before major human trade routes were established (e.g., before the industrial revolution). Conversely, a non-native or exotic species is one that is introduced — intentionally or accidentally — outside of its historical native range. However, for many species, defining the native range is inherently challenging, predominantly due to a lack of rigorous historical records. Only an extremely careful review of available records can accurately delimit native versus non-native ranges (Orlova-Bienkowskaja & Volkovitsh 2018). Moreover, indigenous human knowledge systems and traditional ecological knowledge are often underutilized in these reconstructions of native range (Moller et al. 2004; Fleischman & Briske 2016; Ferreira-Rodriguez et al. 2021). Because of these limitations, many studies accentuate that the dichotomy between native and non-native status should be approached cautiously, particularly when assessing origin (Argueta-Guzmán et al. 2023).

We adopt the following definition of an invasive population as a non-native population that has gone through the stages of invasion defined by Blackburn et al. (2011) and that may cause, or that has the potential to cause, harm to humans through direct competition for resources by imposing health risks or by ecosystem restructuring. The US Department of Agriculture’s National Invasive Species Information Center (USDA NISIC) adopts a similar framework and defines an invasive species as one that is non-native to the ecosystem under consideration and whose introduction causes, or is likely to cause, economic or environmental harm or harm to human health (Executive Order 13112). Blackburn et al. (2011) include in their definition that invasive populations are self-sustaining and capable of range expansion into a new, non-geographically contiguous environment. Furthermore, through the invasion pathway defined by Sorte & Kilman (2016), the establishment of an invasive species is defined as the formation of a reproducing and self-sustaining population without continued human aid after introduction. The distinction between invasive populations and invasive species should also be noted, since some populations of a species may not be self-sustaining or environmentally harmful in a non-native region (Pereyra 2016). Here, we use the term ‘invasive species’ to simply refer to species with known invasive populations.

Projections of biological invasions have rapidly increased and are quite daunting, due in part to increased international trade and commercial activity. For instance,□Levine and□D’Antonio (2003) applied a Michaelis-Menten style model to estimate that 115 new insect species should be expected in the US by 2020. The United States Register of Introduced and Invasive Species (US-RIIS) shows that in 2022, more than 358 new invasive insect species had been recorded in the United States since Levine and D’Antonio’s prediction in 2003 (Simpson et al. 2022). Moreover, once an invasive population becomes established, it is unlikely to be eradicated (Lockwood et al. 2013). Thus, proactive management is required to prevent the establishment of non-native species. Indeed, proactive rather than reactive management is increasingly feasible: early intervention has been shown to offer enormous payoff (Venette et al. 2021). In the context of invasive species management, rapid-response efforts substantially lower control and eradication costs (Koch et al. 2020). Despite the critical need for preemptive measures, there are limited predictive tools to assess the invasive potential of arthropods.

As such, we developed a predictive model that aggregates key life-history and ecological traits associated with highly successful arthropod invaders to estimate invasion potential. To evaluate the model, we selected the invasive honey bee parasite *Tropilaelaps mercedesae*. An emerging brood parasite of honey bees, *T. mercedesae* has generated increasing concern due to its recent range expansion. Importantly, many researchers suggest that its current distribution likely represents only the early stages of a broader global spread (Chantawannakul et al., 2018). The potential expansion of *T. mercedesae* is particularly concerning because the western honey bee (*Apis mellifera*) underpins a substantial global pollination industry and food production system. In the United States alone, honey bees are estimated to contribute between $14.6 billion and over $18 billion annually, depending on pollination-service and honey-production estimates (Calderone, 2012; USDA, 2025). Globally, honey bees are integral to pollination in both agricultural and natural ecosystems (Hung et al., 2018; Requier et al., 2019; Reilly et al., 2024). This vast global enterprise is further supported by extensive efforts to reduce colony losses and maintain bee health (Ziegler et al., 2021). Given the economic, ecological, and cultural importance of *A. mellifera*, the continued expansion of *T. mercedesae* poses substantial risks. As invasion pressure increases, the ability to rapidly detect and accurately diagnose emerging parasites becomes critical for early containment and mitigation. Therefore, identifying high-risk invasive species through predictive modeling must be coupled with the development of sensitive, field-deployable diagnostic tools to safeguard honey bee health and agricultural sustainability.

Currently, apiculture is challenged by severe colony loss, driven by pesticide exposure, nutritional stress, parasites and pathogens, and climate change (Smith et al. 2013; Paudel et al. 2015; Hristov et al., 2021). These stressors are all exacerbated by the mite *Varroa destructor* (Class: Arachnida, Order: Mesostigmata), a cosmopolitan invasive arthropod and well-known driver of honey bee colony loss (Noël et al. 2020; Sankovitz et al. 2025). Originally discovered in 1904 on eastern honey bees (*Apis cerana*) hosts, *V. destructor* is now known to parasitize the brood of *A. mellifera*, feed on adult and juvenile bee fat-body tissue, and act as a vector for multiple viruses — thereby weakening bees, reducing longevity, and decreasing colony survival (Ramsey et al. 2018, 2019; Traynor et al. 2020; Sonenshine et al. 2022). Although treatments have been developed to curb mite populations, *V. destructor* has quickly developed tolerance and resistance to many of these treatments (Mitton et al. 2022). The rapid spread of *V. destructor* after acquiring *A. mellifera* as a host, coupled with repeated failures to limit its expansion, underscores the critical importance of predicting biological invasions before they spread, establish, and cause harm (Blackburn et al. 2011).

Given the damage *A. mellifera* has incurred from *V. destructor*, there is increasing concern about *Tropilaelaps* mites, specifically *T. mercedesae* (Chantawannakul et al. 2018; Ramsey 2021). The *Tropilaelaps* genus was discovered in 1961 in the Philippines (Delfinado and Baker). Current evidence indicates that *Tropilaelaps* mites shifted from wild honey bee hosts—*Apis dorsata*, *Apis laboriosa*, and *Apis breviligula*—to *A. mellifera* in Southeast Asia (Forsgren et al. 2009; de Guzman et al. 2017). These mites share key life-history traits with *V. destructor* mites — for example, reproduction within capped brood cells and parasitism of developing bees — but they also differ in ways that may increase threat. They inflict more wounds when feeding, have a short or possibly nonexistent non-reproductive phase, and seem to reproduce more rapidly (de Guzman et al. 2017; Phokasem et al. 2019; Ling et al. 2023).

More concerning, a growing body of research indicates that *Tropilaelaps* mites may act as disease vectors, like their *Varroa* mite counterparts (Dainat et al. 2008; Forsgren et al. 2009; Khongphinitbunjong et al. 2015, 2016; Wu et al. 2017; Chanpanitkitchote et al. 2018; Truong et al. 2023). Currently, *Tropilaelaps* mite populations are rapidly expanding their range across Asia and into Europe, posing a major emerging threat to beekeeping and pollination services (Janshia et al. 2024; Joharchi & Stolbova 2024; Mohamadzade Namin et al. 2024; Brandorf et al. 2025). Despite efforts to quantify and track the presence and spread of *Tropilaelaps*, little work has been done to describe the mite’s life history and ecology, highlighting a need for further prophylactic research to mitigate invasion events. Highlighting the potential risk posed by *Tropilaelaps* mites before they arrive in the Americas and spread across Europe may facilitate crucial early detection, surveillance, and policy mobilization. Moreover, little research has been conducted to investigate *T. mercedesae*’s true invasive potential despite its rapid spread into non-native regions. Thus, *Tropilaelaps* mites provide an excellent case study for applying existing data on invasive arthropods to an emerging non-native species.

**Figure 1.**
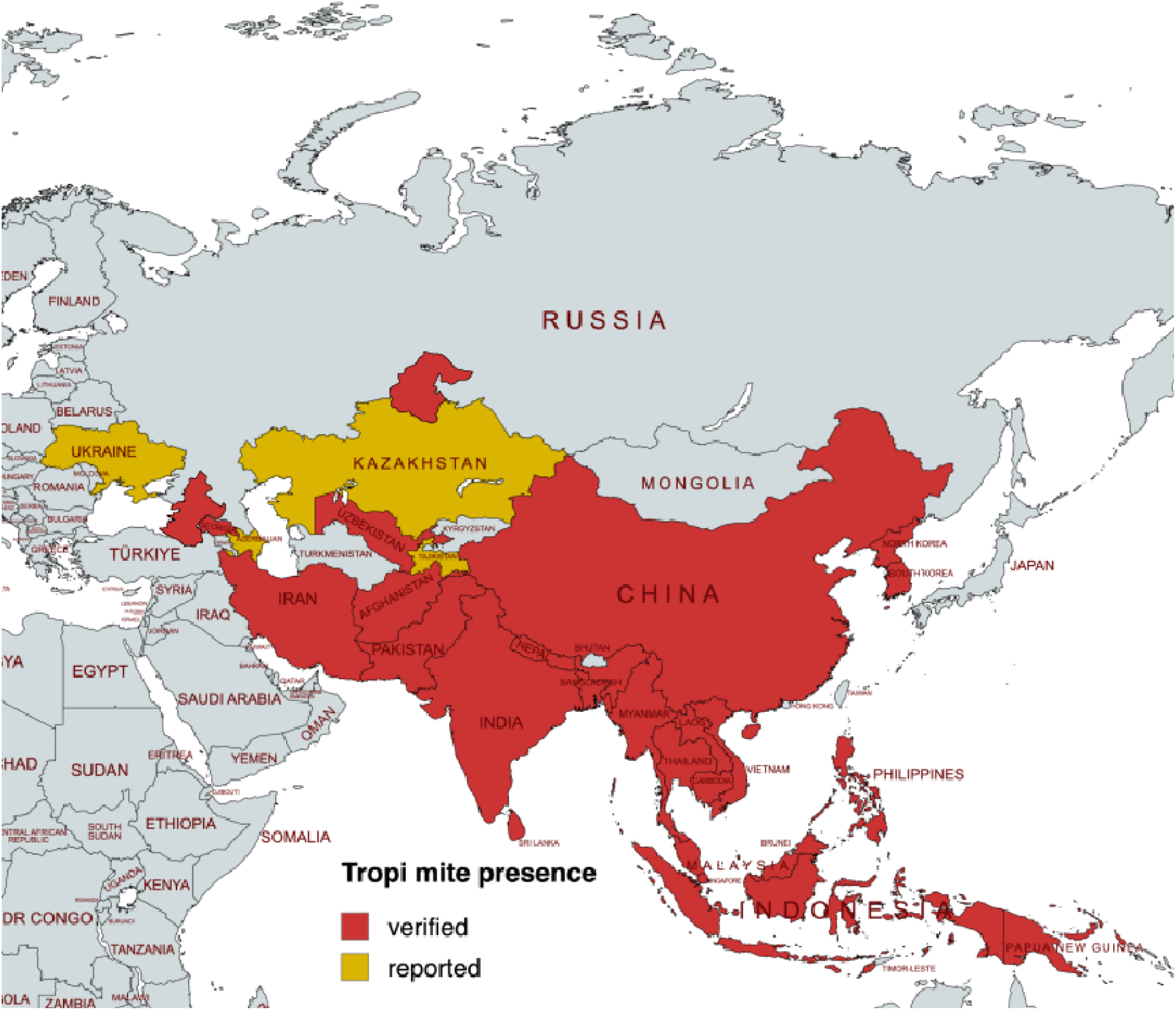
Map of *Tropilaelaps mercedesae* mites reported and verified presence. Red fill indicates verified presence of *T. mercedesae*. Yellow fill indicates reported but unverified presence of *T. mercedesae.* See supplementary information for references. Due to existing surveillance gaps, there are no current reports of *T. mercedesae* in Bhutan, Turkmenistan, and Kyrgyzstan, but they are likely present due to reports on surrounding borders. Citations for each report can be found in Supplementary File 1.

To identify which biological traits and species-specific environmental constraints best predict invasion success in arthropods, we compiled a comprehensive trait-based dataset of invasive species and used it to construct a comparative statistical model of invasibility.

Following model selection, diagnostics, and validation, we applied the top-performing model to *T. mercedesae* to estimate its capacity to establish invasive populations relative to other known invasive arthropods. A trait-based comparative framework enables evaluation of *Tropilaelaps* mites’ potential to become a cosmopolitan invasive species akin to *Varroa*, providing a preemptive, data-driven assessment of the invasion risk posed by this emerging threat to apiculture.

## Methods

### Dataset Curation

To quantify the traits associated with invasive species, we first compiled known invasive arthropod species from the Global Invasive Species Database (GISD 2025). We limited our scope to the phylum Arthropoda given that it has substantially more invasive species than all other animal phyla (IPBES 2023; Seebens 2025). This is likely attributable to their short reproductive cycles, spatial ubiquity, and diverse trophic niche occupancy. Once all arthropod entries were amassed from GISD, all authors reviewed the existing dataset and additional invasive arthropod entries were added to create a more robust, encompassing dataset. These additions were made to capture underrepresented ecological niches, taxonomic groups, and invasion strategies, thereby improving the breadth and coverage of the dataset. Following review of the dataset, we added notable missing arthropod species, including but not limited to *Anopheles stephensi* (Asian malaria mosquito)*, Blattella germanica* (German cockroach), *Bombus terrestris* (buff-tailed bumblebee), *Helicoverpa armigera* (corn earworm), *Leptinotarsa decemlineata* (Colorado potato beetle), *Loxosceles rufescens* (Mediterranean recluse), *Lucilia cuprina* (Australian sheep blowfly), and *Periplaneta americana* (American cockroach). This yielded a dataset containing 127 invasive arthropods.

A literature review of all entries yielded information on the current distribution of these species, the associated environmental variables of their distributions, and the life history traits of these entries (Whitney & Gabler 2008; Weir & Salice 2011; Hodgins et al., 2018). All collected data for each arthropod in the dataset can be found in Supplementary File 2. This led to the identification of potential predictor traits for quantifying the adaptive niche of known invasive species (Table 1). All parameters were defined for each species using primary literature or publicly available data sets. Data sets included the National Oceanic and Atmospheric Administration (NOAA) Climate Assessment Database for 1991-2020 (National Weather Service 2025), the Centre for Agriculture and Bioscience International (CABI) database confirmed reports, and, if needed, certified Global Biodiversity Information Facility (GBIF) records. For additional information on each parameter’s subdivided levels or criteria, see Supplementary File 2. To reduce bias while maintaining consistency and accuracy, the dataset was independently curated by three researchers, and all entries were subsequently reviewed by one researcher of the three researchers. Species with unreliable data, missing parameters, or that did not meet the standards of the dataset were removed. This included species bred in captivity with no native range and released, or species with highly disputed native ranges. All removed species are listed in Table 2. Our final dataset contained 118 invasive arthropods.

**Table 1.**
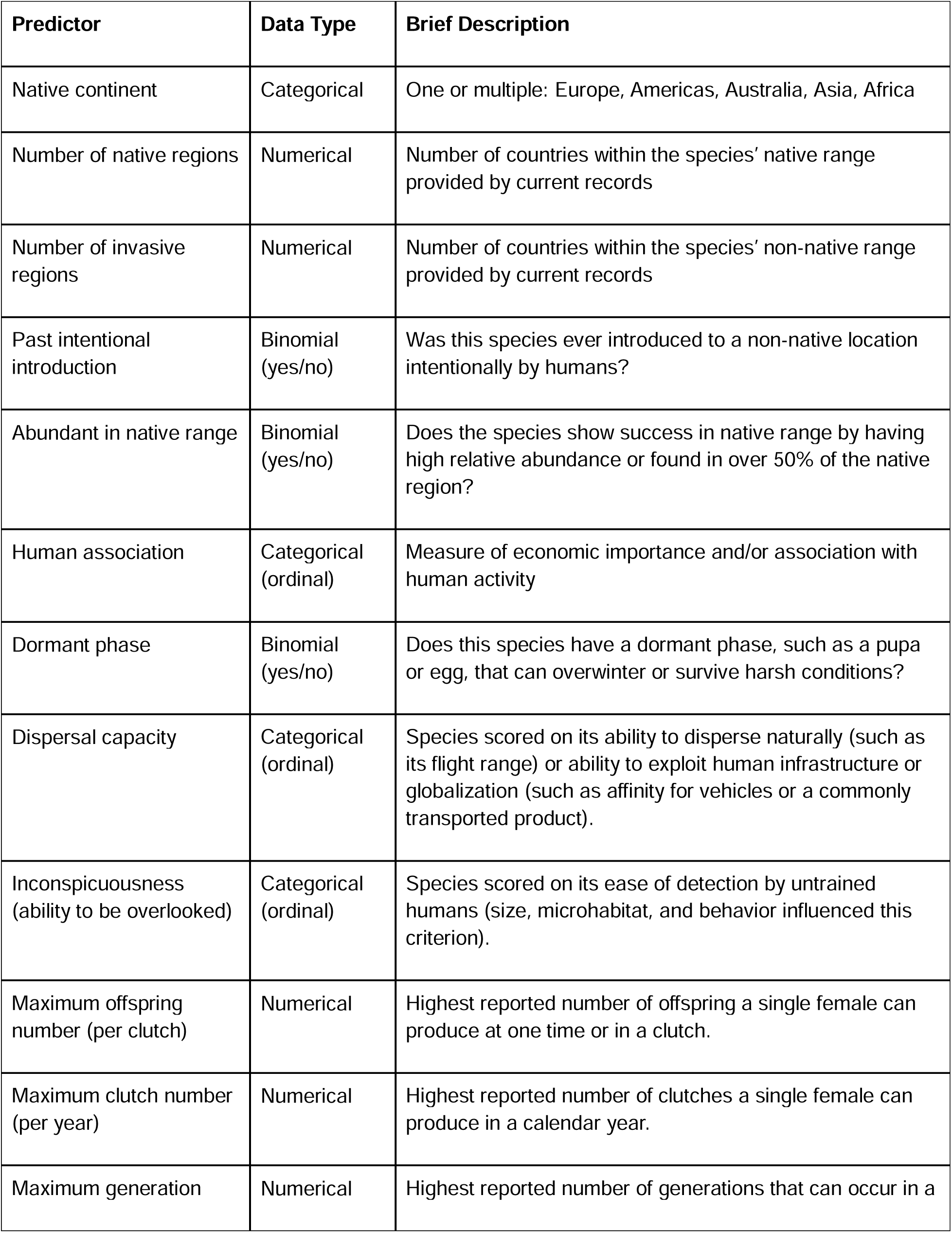
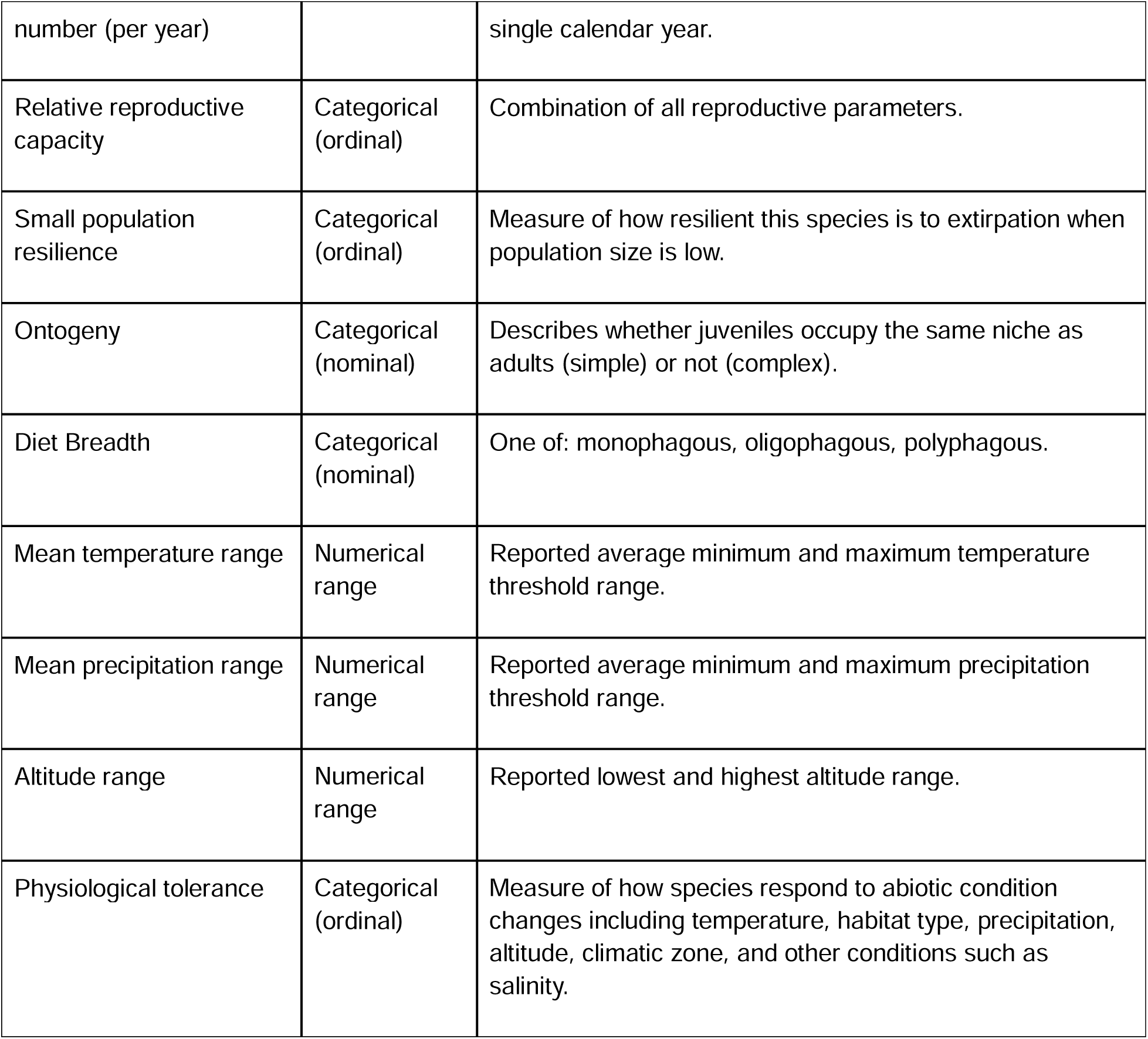
Data collected from literature to quantify the adaptive niche of known invasive species. See Supplementary File xx for more information on each trait and how it was defined.

**Table 2.** Arthropod species removed from the dataset with respective reasons.

| Species | Common Name | Removal Reason |
| --- | --- | --- |
| *Dysdera crocata* | woodlouse spider | Insufficient peer-reviewed data |
| *Nylanderia fulva* | raspberry crazy ant | Insufficient peer-reviewed data |
| *Hoplochelus marginalis* | white sugar cane worm | Insufficient peer-reviewed data |
| *Ornithonyssus bacoti* | tropical rat mite | Cosmopolitan distribution;<br>no distinct native range |
| *Oryctes rhinoceros* | coconut rhinoceros beetle | Insufficient peer-reviewed data |
| *Procambarus virginalis* | marbled crayfish | No true native range, bred in captivity |
| *Sitophilus oryzae* | rice weevil | Insufficient peer-reviewed data |
| *Trogoderma variabile* | warehouse beetle | Insufficient peer-reviewed data |
| *Vespa crabro* | European hornet | Insufficient peer-reviewed data |

### Trait Selection Rationale

Findings broadly in the field of invasion ecology led to the selection of traits which have been shown consistently across multiple studies to contribute substantially to the success of would-be invaders. These traits can contribute at any or all of the three stages of invasion: incursion/introduction, establishment, and spread. **High relative abundance** in an organism’s native range suggests that an organism is adept at living in a broad range of habitats (Jeschke & Strayer 2005). The variable suite of adaptations conferring this capacity have been shown to render the putative invader a markedly successful competitor for resources outside of its established geographic range as well (Jeschke & Strayer 2005; Pyšek et al. 2009). Close **human association** is a key predictor of invasibility with numerous examples in fish, livestock, ornamental plants, and insects such as honey bees (Garcia-Berthou 2007; Jeshke & Strayer 2006; Lockwood et. al. 2013; Russo et. al. 2021). Human-affiliated organisms show increased likelihood of successful introduction, establishment, and spread as humans serve as transport vectors and cultivators often substantially increasing the propagule pressure at the point of incursion (Garcia-Berthou 2007; Jeshke & Strayer 2006; Lockwood et. al. 2013). Potting soil, clothing, attachment to ships/automobiles, ballast water, etc. have caused humans to unknowingly transport countless arthropods from yellow-legged hornets (*Vespa velutina*), red imported fire ants (*Solenopsis invicta*), tropical bed bugs (*Cimex hemipterus*), brown marmorated stinkbugs (*Halyomorpha halys*) and chinese mitten crabs (*Eriocheir sinensis*) (Cohen and Carlton 1997; Hoebeke and Carter 2003; Ross and Shoemaker 2008; Campbell et al. 2016; Keeling et al. 2017).

Additionally, a would-be invasive organism has no guarantee it will be introduced into an environment when conditions are favorable and as such can benefit substantially from a physiological **dormant phase** (Redwood et al. 2019). Dormancy (expressed as torpor, hibernation, or heightened resilience in an inactive lifestage like eggs or pupae) buffer the population against harsh climates and unfavorable seasons (Redwood et al. 2019; Pyšek et al. 2009). It further synergizes with human association and **inconspicuousness** to allow organisms to easily be relocated (e.g. *V. velutina* queens dormant in pottery or small *H. halys* egg masses on ornamental plants) (Hoebeke and Carter 2003; Keeling et al. 2017).

**Inconspicuous** organisms have an advantage in all stages of invasion allowing for easier introduction to new ecosystems via human assistance and infrequent detection thereafter (Lockwood et al. 2013; Morais and Reichard 2018). Similarly, broad environmental tolerance (i.e. **temperature tolerance, precipitation tolerance, elevational resilience,** and generally **high physiological tolerance**) allows a putative invader to persist even in environments considered prohibitively hostile to other species (Pyšek et al. 2009; Pyšek et al. 2011; Pauchard et al. 2009; Higgins & Richardson 2014).

Organisms with **high dispersal capacity** are especially adept in the spread stage of ecosystem invasion because they routinely move large distances already as a natural part of their lifecycle (Phillips et al. 2006; Pysek et al. 2009). Studies of invasibility have consistently shown that organisms with high fecundity are also highly adept invaders as the process tends to favor the *r*-selected strategy of treating survival like a numbers game (Tschinkel 2006; Lockwood et al. 2013; Kistner et al. 2017; Liu et al. 2020; Seebens et al. 2025). Each offspring represents an opportunity for the right set of circumstances to align which allow the organism to escape stochastic challenges that would otherwise lead to the extirpation of a smaller population (Lockwood et al. 2013). We’ve divided the category of high fecundity between those arthropods with high offspring output in a single reproductive event (**high maximum offspring**), those with many reproductive events per year (**high maximum clutch**), and those with multiple generations per year (**high maximum generation number**) as each contributes differently to invasion capacity (Moyle and Light 1996; Liu et al. 2020; Currylow et al. 2023). Moreover, organisms with **high small population resilience** can resist extirpation even when population size is low, such as the ability to reproduce through parthenogenesis or resistance to inbreeding (Baker 1955; Elam 2007; Gutenkunst 2018). Organisms such as *V. destructor* and *V. velutina* use gravid females as dispersal units such that a single foundress can effectively establish a successful population (Keeling et al. 2017; Rosenkranz 2010).

We’ve long understood the benefit of immature and adult organisms of the same species focusing their attention on different food sources, but it is becoming clearer that this life history strategy can be especially helpful in the process of ecosystem invasion (Kupferberg 1997; Simberloff and Rejmanek 2011). Sorensen and Bergstedt (2011) were able to show that the **complex ontogeny** of filter-feeding juvenile sea lampreys and the parasitic adults promoted their success in all three stages of the ecosystem invasion process. Additionally, **broad diet breadth** not only reduces competition for food resources but further allows ecosystem invaders to thrive in additional locations that would be unfavorable to a specialist lacking its sole food source (Case & Gilpin 1974; Lockwood et al. 2013).

We further assessed the point of origin for the species by focusing on its **native continent** and the **number of native regions** as invasive species follow a pattern of invasion from certain regions which are more inclined to serve as donors rather than recipients (e.g. Southeast Asia as donor and the US at recipient) (Pysek et al. 2009; van Kleunen et al. 2016). However, there is no stronger predictor of whether a species will become invasive in a region than whether it is already invasive in another part of the world (Lockwood et al. 2013; Seebens et al. 2025). As such, we additionally included in our model the **number of invasive regions** of a putative invader and examined **past intentional introduction** to determine *how* a species reached novel ecosystems prior.

### Model Selection

Model selection was conducted to identify traits associated with arthropod invasibility.

We used the number of invasive regions as the response variable, with a higher count indicating a more invasive species or higher invasive potential. We treat this variable as a proxy for each arthropod species’ inherent invasive ability.

Predictor variables were first assessed for correlations, and for highly correlated variables, only one was retained. Maximum number of generations per year was selected over maximum offspring per clutch, maximum clutches per year, relative reproductive capacity, and the ability to have a dormant phase. Physiological tolerance was represented by altitude, temperature, and precipitation ranges and therefore was not included as a separate variable. Human association was retained over dispersal capacity, and ability to enter a dormant phase was retained over species conspicuousness. Abundance in the native range was excluded because 110 of 118 invasive species were classified as abundant. This filtering process resulted in seven predictor variables: human association, ability to enter a dormant phase, maximum number of generations per year, diet breadth, mean temperature range, precipitation range, and altitude range.

Ordinal variables were rebinned to evenly distribute entries across categories. For example, the human association variable was initially categorized into five ordinal levels: *very low, low, moderate, high,* and *very high*. Owing to sparse arthropod representation in the *very low* and *very high* categories, these levels were consolidated with adjacent categories, resulting in three ordinal levels: *low, moderate,* and *high*. In addition, numerical predictors were scaled as needed to improve model performance, maximize interoperability, and reduce potential collinearity (Chatterjee & Hadi, 2015). Specifically, temperature range, altitude range, and precipitation range were scaled and/or centered.

A negative binomial generalized linear mixed model (GLMM) was implemented using the R package glmmTMB (Brooks et al. 2017), given the distribution of the number of invasive regions in the dataset (variance > mean). All combinations of predictor variables were tested and compared to the full model, including all predictors. Random effects for native continent and taxonomic class were included as random intercepts to account for phylogenetic and human trade biases. Models were evaluated using the Akaike Information Criterion (AIC), and the top models were compared using likelihood-ratio tests (ANOVA). Subsequently, significant interactions between predictor variables were tested. Interactions were removed if non-significant.

Model selection was repeated with a subset of our dataset (n = 106) excluding species intentionally introduced by humans to a non-native location (n = 12). This was done to account for population introductions that were not a direct function of the species’ trait ensemble. Finally, the most significant and best-performing model was evaluated using the DHARMa package (Hartig 2025) to assess zero-inflation, fitted and residual alignment, and possible covariance errors. As an additional evaluation of overall model performance, *Halyomorpha halys* (brown marmorated stink bug), *Popillia japonica* (Japanese beetle), *Pectinophora gossypiella* (pink bollworm), and *Amblyomma variegatum* (tropical bont tick) were intentionally excluded from the curated dataset during model construction. Model performance was then assessed by comparing the predicted number of invasive regions for *H*. *halys*, *P*. *japonica*, *P*. *gossypiella*, and *A*. *variegatum* with the empirically verified number of regions within its current invasive distribution.

### Model Application to *Tropilaelaps* Mites

Following diagnostic assessment, the top-performing model was used to predict the number of invasive regions for *Tropilaelaps mercedesae*. This species’s predicted number of invasive ranges was then ranked on our list of known invasive arthropods based on the number of observed invasive ranges for comparison. *T. mercedesae* data for each selected predictor variable were acquired from primary literature and from personal unpublished data collected from international research trips; the data are viewable in the supplementary material.

## Results

### Model

After all model selection and interactions were tested, the top-performing model was selected including human association, small population resilience, diet breadth, maximum annual generations, temperature range, altitude range, and the interaction between human association and temperature range as significant predictors for the number of invasive regions of an arthropod. To reduce backdoor confounding, the same dataset (excluding any species that was intentionally introduced at least once in the historical record) was used for model selection and the same top-performing model was selected. Our top-performing model was then verified through the DHARMa test for zero inflation (Figure S1), residual vs. fitted output (Figure S2), and covariance (Figure S3). All model diagnostics indicated that the selected model satisfied the underlying assumptions, with no significant issues related to the covariance structure, goodness-of-fit, or zero inflation. In addition, based on the current literature, our model predicted that *Halyomorpha halys* would be invasive in 79 regions; *H. halys* has been verified as invasive in 71 regions. *Popillia japonica* is confirmed invasive in 50 regions and was predicted to be invasive in 60. *Pectinophora gossypiella* invasion potential was estimated to be 81 regions but is currently established in 107 regions. Finally, *Amblyomma variegatum* has been established in 23 regions primarily throughout the Caribbean but was estimated to be able to invade 28 regions. Summary of these predictions can be found in Table 3.

**Table 3.** Intentionally excluded arthropods from the dataset used to build our model, their predicted invasive potential (predicted number of invasive regions), and their current confirmed number of invasive regions.

| Species | Order | Predicted Invasive Potential | Confirmed Invasive Regions |
| --- | --- | --- | --- |
| *Halyomorpha halys* | Hemiptera | 79 | 71 |
| *Popillia japonica* | Coleoptera | 60 | 50 |
| *Pectinophora gossypiella* | Lepidoptera | 81 | 107 |
| *Amblyomma variegatum* | Ixodida | 28 | 23 |

### Top performing model

*# of introduced (invasive) regions ∼ human association + small population resilience + diet breadth + maximum annual generations + temperature range + altitude range + human association: temperature range + (1| class/order) + (1| native continent)*

Our model showed that the intercept for all arthropods was significant, with a log count predicted invasive regions of 2.3138 ± 0.3686. An arthropod with moderate human association significantly increased the log count of predicted invasive regions by 0.5329 ± 0.2354 compared to arthropods with low human association. An arthropod with high human association significantly increased the log count of predicted invasive regions by 0.9585 ± 0.2233 compared to arthropods with low human association. An arthropod with a polyphagic diet significantly increased the log count of predicted invasive regions by 0.8947 ± 0.0157 compared to arthropods with a monophagic diet. An arthropod that is capable of having 11-28 generations in a single calendar year significantly increased the log count of predicted invasive regions by 0.6666 ± 0.2981 compared to arthropods that only had a single generation per calendar year.

However, an arthropod that had continuous generations significantly decreased the log count of predicted invasive regions by 0.5581 ± 0.2713 compared to arthropods that only had a single generation per calendar year. Most eusocial organisms, such as ants, lack distinct generations, as colonies continually produce offspring, especially taxa that exhibit polygyny or supercolony formation. For each one-meter increase in altitude range, the log count of predicted invasive regions increased by 0.3588 ± 0.1088 per arthropod. A significant interaction between high human association and temperature range was also observed; for arthropods with high human association, each 1°C increase in temperature range was associated with an increase of 0.0306 ± 0.0150 in the log count of predicted invasive regions per arthropod. Model outputs are presented in Tables 4 and 5.

**Table 4.** All outputs (estimate, standard errors, and p-values) for top performing model.

| Category | Subcategory | Estimate | Standard Error | p-value |
| --- | --- | --- | --- | --- |
| **Intercept** |  | **2.3138** | **0.3686** | **<0.0001\*\*\*** |
| **Human Association** | Low (<u>reference level</u>) |  |  |  |
|  | Moderate | **0.5329** | **0.2354** | **<0.05\*** |
|  | High | **0.9585** | **0.2233** | **<0.0001\*\*\*** |
| **Small Population Resilience** | Low (<u>reference level</u>) |  |  |  |
|  | High | 0.2995 | 0.2026 | >0.05 |
|  | Very High | **0.9276** | **0.2437** | **<0.0001\*\*\*** |
| **Diet Breadth** | Monophagic (<u>reference level</u>) |  |  |  |
|  | Oligophagic | 0.05304 | 0.3031 | >0.05 |
|  | Polyphagic | **0.8947** | **0.0157** | **<0.001\*\*** |
| **Max Annual Generations** | 1 (<u>reference level</u>) |  |  |  |
|  | 2-10 | 0.0048 | 0.2346 | >0.05 |
|  | 11-28 | **0.6666** | **0.2981** | **<0.05\*** |
|  | 29-54 | -0.3156 | 0.4934 | >0.05 |
|  | Continuous | **-0.5581** | **0.2713** | **<0.05\*** |
| **Temperature Range** | (numerical) | 0.0041 | 0.0091 | 0.05 |
| **Altitude Range** | (numerical) | **0.3588** | **0.1088** | **<0.001\*\*** |
| **Human Association x** | Low (<u>reference level</u>) x |  |  |  |

| Temperature Range | Temperature Range |  |  |  |
| --- | --- | --- | --- | --- |
|  | Moderate x Temperature Range | 0.0147 | 0.01929 | 0.05 |
|  | High x Temperature Range | **0.0306** | **0.0150** | **= 0.05\*** |

\* Denotes significant p-value equal or less than 0.05, but greater than 0.001.
\*\* Denotes significant p-value equal or less than 0.001, but greater than 0.0001.
\*\*\* Denotes significant p-value equal or less than 0.0001.

**Table 5.** Significant outputs (estimate, standard errors, and p-values) for the top performing model.

| **Category** | **Subcategory** | **Estimate** | **Standard Error** | **p-value** |
| --- | --- | --- | --- | --- |
| **Intercept** |  | **2.3138** | **0.3686** | **<0.0001\*\*\*** |
| **Human Association** | **Moderate** | **0.5329** | **0.2354** | **<0.05\*** |
| **Human Association** | **High** | **0.9585** | **0.2233** | **<0.0001\*\*\*** |
| **Small Population Resilience** | **Very High** | **0.9276** | **0.2437** | **<0.0001\*\*\*** |
| **Diet Breadth** | **Polyphagic** | **0.8947** | **0.0157** | **<0.001\*\*** |
| **Max Annual Generations** | **11-28** | **0.6666** | **0.2981** | **<0.05\*** |
| **Max Annual Generations** | **Continuous** | **-0.5581** | **0.2713** | **<0.05\*** |
| **Altitude Range** | **(numerical)** | **0.3588** | **0.1088** | **<0.001\*\*** |
| **Human Association x Temperature Range** | **High x Temperature Range** | **0.0306** | **0.0150** | **= 0.05\*** |

\* Denotes significant p-value equal or less than 0.05, but greater than 0.001.
\*\* Denotes significant p-value equal or less than 0.001, but greater than 0.0001.
\*\*\* Denotes significant p-value equal or less than 0.0001.

### Model Application to *Tropilaelaps* Mites

Once the model passed diagnostic tests, data for *Tropilaelapes mercedesae* were used to predict the number of invasive regions for the emerging invasive species. When the number of invasive regions was simulated using the model for all selected species,*T. mercedesae* was the 12^th^ most adept invader with 160 predicted invasive regions (Table 6). This places *T. mercedesae* in the top 10% of invasive arthropods (or the 91^st^ percentile in this dataset for invasive potential). If placed directly into the original dataset (Table 7), *T. mercedesae* would rank as the seventh most adept invasive species among 119 invasive arthropods worldwide.

**Table 6.** Top 20 invasive species from the top performing model and predicted number of invasive regions.

| Species | Predicted Number of Invasive Regions |
| --- | --- |
| *Harmonia axyridis* | 852.7112868 |
| *Phenacoccus solenopsis* | 298.3440269 |
| *Periplaneta americana* | 255.4988565 |
| *Culex quinquefasciatus* | 218.392631 |
| *Rhyzopertha dominica* | 216.9362081 |
| *Calliphora vicina* | 211.5270479 |
| *Leptinotarsa decemlineata* | 209.7795037 |
| *Aedes albopictus* | 172.7427391 |
| *Varroa destructor* | 170.5395628 |
| *Ceratitis capitata* | 169.7168374 |
| ***Tropilaelaps mercedesae*** | **160.4164577** |
| *Aedes aegypti* | 135.1734668 |
| *Bemisia tabaci* | 128.2250641 |
| *Drosophila suzukii* | 124.7335472 |
| *Loxosceles rufescens* | 124.3224378 |
| *Blattella germanica* | 116.7216949 |
| *Pheidole megacephala* | 114.6187091 |
| *Helicoverpa armigera* | 112.2569197 |
| *Maconellicoccus hirsutus* | 110.1466481 |
| *Spodoptera frugiperda* | 93.12108989 |

**Table 7.** Top 20 invasive species from literature review and number of invasive regions, excluding *Tropilaelapes mercedesae*.

| Species | Reported Number of Invasive Regions |
| --- | --- |
| *Periplaneta americana* | 960 |
| *Leptinotarsa decemlineata* | 822 |
| *Culex quinquefasciatus* | 487 |
| *Bemisia tabaci* | 280 |
| *Loxosceles rufescens* | 206 |
| *Icerya purchasi* | 181 |
| *Paratrechina longicornis* | 156 |
| *Varroa destructor* | 135 |
| *Harmonia axyridis* | 129 |
| *Blattella germanica* | 126 |
| *Phenacoccus solenopsis* | 119 |
| *Lucilia cuprina* | 114 |
| *Tapinoma melanocephalum* | 114 |
| *Plutella xylostella* | 112 |
| *Drosophila suzukii* | 110 |
| *Maconellicoccus hirsutus* | 106 |
| *Aedes aegypti* | 104 |
| *Aedes albopictus* | 95 |
| *Porcellio scaber* | 92 |
| *Insignorthezia insignis* (*Orthezia insignis*) | 92 |

Additionally, the simulated ranking of invasive species by predicted number of invasive regions closely matched the ranking from observed invasive range values.

## Discussion

Our results demonstrate that a small set of biological traits and species-specific environmental constraints can reliably predict the global invasibility of arthropod taxa. Specifically, human association, small-population resilience, number of annual generations, and broad environmental tolerance emerged as the key predictors of invasion success across diverse arthropod lineages. These traits are consistent with long-standing invasion hypotheses, including the roles of propagule pressure, ecological generalism, and physiological flexibility in increasing the likelihood that introduced populations will overcome demographic and environmental barriers (Levine & D’Antonio 2003; Blackburn et al. 2011). The consistency of these findings across taxonomic groups suggests that trait-based models can complement pathway-and trade-based risk assessments, offering a biologically grounded lens for forecasting future invaders before their arrival (Fournier et al. 2019). However, certain traits, such as native range abundance, were not included in the model due to high homogeneity among species. Though this was not a useful predictor of invasibility, it is a trait conserved in invasive arthropods. This trend has been noted in other invasive species but is not universal Parker et al. 2013; Hui et al. 2023). This observation may be due to high native abundance correlating with ecosystem generalism, increasing an individual’s odds of being transported by humans or other vectors, being linked to increased genetic diversity, or to the biology of arthropod reproduction strategies and rapid development. While many invasive species are common or abundant in their native ranges, high native range abundance is neither necessary nor sufficient for invasiveness (Hansen et al. 2013; Parker et al. 2013).

Human association was one of the strongest predictors in our best-fit model, reinforcing the longstanding observation that species frequently transported or provisioned by humans often have elevated invasion probabilities. Human movement increases propagule pressure (Lockwood et al. 2013; Simberloff 2009), facilitates repeated introductions (McDermott & Finoff 2016; Bertelsmeier 2018), and provides disturbed or synanthropic habitats that can buffer small population sizes during early establishment (Wilson et al. 2009; Guo et al. 2018). Likewise, high reproductive potential – indexed here by maximum annual generations – was associated with greater invasibility, which aligns with theory predicting that fast life histories increase the chance of surviving stochastic population bottlenecks during introduction and spread (Allen et al. 2017). Broad physiological tolerance, reflected by wide temperature and altitude ranges, supports the idea that environmental flexibility enables invaders to colonize climatically diverse regions and expand beyond the initial point of establishment.

In order to assess overall model performance, we purposefully excluded a diverse group of arthropods, including a hemipteran, lepidopteran, coleopteran, and ixodid. These species were selected because they exhibit diverse invasion strategies and variable management efficacy. Overall, our top-performing model closely predicted the number of established invasive regions for each species. *Halyomorpha halys* is an important agricultural and household pest in North America. Our model predicted that this species would occur in 79 invasive regions, and it is currently confirmed in 71 regions (Dioli et al. 2016; Hamilton et al. 2018; Musolin et al. 2018; Berteloot et al. 2024). However, many researchers predict that *H. halys* will continue to spread, particularly into environmentally suitable areas such as northern Europe, northeastern North America, southern Australia, and the North Island of New Zealand (Zhu et al. 2012; Haye et al. 2015). Similarly, our model slightly overestimated the confirmed range of *Popillia japonica*, predicting invasion in 60 regions compared to 50 confirmed regions (Kistner-Thomas 2019; Strangi et al. 2024). Given the documented range expansion of this species over the past 100 years, additional spread is likely (Della Rocca & Milanesi 2022; Barré & Uilenberg 2010; Pulighe et al. 2025). *Amblyomma variegatum* is currently established in 23 regions, while our model estimated invasion potential in 28 regions (Phillip 2000; Beati et al. 2012). Although this species has been detected in other regions without establishing populations, concerns remain regarding potential range expansion into Europe and North America via avian or livestock mediated dispersal (Solomon & Kaaya 1998; Estrada-Peña et al. 2007; Pascucci et al. 2007). The model underestimated the invasive range of *Pectinophora gossypiella*, which is currently established in 107 regions (excluding areas where populations have been eradicated following the introduction of Bt cotton), but was predicted to occur in 81 regions (Liu et al. 2009; Staten & Walters 2021; Bhute et al. 2023; Matheson et al. 2023). Collectively, these results demonstrate the model’s ability to estimate invasion potential for well documented invasive species and emphasize that the model predicts overall invasion potential rather than the true number of established regions. This provides strong evidence for the effectiveness and utility of predictive trait-based modeling in assessing arthropod invasion risk.

Applying this model to *Tropilaelaps mercedesae* revealed that the mite exhibits traits most associated with invasion success. Most importantly, *Tropilaelaps mercedesae* is strongly associated with human activity via apiculture. Humans transport honey bees and agricultural materials across the globe regularly, along with *T. mercedesae* parasitizing them. And like *V. destructor*, *T. mercedesae* can produce a new reproductive generation in a matter of days, helping it survive stochastic local extinction events and increasing adaptive speed. Although as a parasite *T. mercedesae* can only survive in the highly controlled microhabitat of a bee hive, the global presence of bee hives and the human maintenance of these hives allows the mite access to a huge range of climates and conditions. Our results show that *T. mercedesae* is amongst the highest-risk taxa in our dataset, indicating substantial potential to establish invasive populations if introduced beyond its current range. This finding is notable given that *T. mercedesae* ranked in invasiveness only slightly below *Varroa destructor* mites, one of the most economically disruptive arthropod invaders in modern agriculture (Traynor et al. 2020). The model therefore reinforces existing biological concerns about *Tropilaelaps* mites while providing a comparative, quantitative context: its combination of rapid reproduction and close human association via apiculture places it among the arthropods most likely to spread globally in coming years.

For apiculture, these results underscore the importance of proactive measures. While *Tropilaelaps* mites have not yet achieved the global distribution of *V. destructor*, our findings suggest that prevention, surveillance, and regulation of managed honey bee movement will be substantially more effective than eradication or suppression after establishment. The history of *V. destructor* mites demonstrates how rapidly an undetected parasite can reshape beekeeping, pollination services, and agricultural economies (Peck 2021). Because *Tropilaelaps* mites share core life-history traits with *V. destructor* mites but possess biological characteristics that may accelerate colony-level damage, their invasion potential warrants serious consideration in biosecurity planning. Notably, this necessitates the urgent need for standardized monitoring and detection methods for the *Tropilaelaps* genus.

However, many gaps remain in our understanding of *Tropilaelaps* biology, ecology, and spread; these results underscore the importance of addressing this knowledge gap. Key life-history parameters—including reproductive rates, survivorship, and population growth thresholds under field conditions—are poorly quantified, particularly across environmental gradients and management contexts (Anderson & Morgan 2007; de Guzman et al. 2017). In addition, the current literature is characterized by numerous unsubstantiated reports of various *Tropilaelaps* species occurring on a variety of hosts, including *Apis cerana*, *Apis florea*, *Bombus* spp., Rodentia spp., and *Xylocopa* spp., as well as in regions such as Kenya and Tasmania (Delfinado-Baker 1982; Laigo and Morse 1968; Kapil and Aggarwal 1987; Bhasker and Putatunda 1989; Kumar et al. 1993; Abrol and Putatunda 1995; Abrol and Putatunda 1996; Ellis and Munn 2005; EFSA Panel on Animal Health and Welfare [AHAW] 2013). Transmission ecology also remains unresolved, and the relative importance of human-mediated colony transport has not been empirically assessed. Finally, although *T. mercedesae* has been reported in a variety of climatic zones and their host colonies shelter them from harsh conditions, the environmental limits of *Tropilaelaps* mite establishment, including sensitivity to climate, seasonality, and brood availability, remain poorly defined, hindering predictive spread modeling and biosecurity prioritization. Together, these gaps emphasize that proactive investment in integrative research—spanning life-history biology, host–parasite interactions, and movement networks—is essential if *Tropilaelaps* mites are to be detected, predicted, and contained before they replicate the transformative impacts of *Varroa* mites.

While the implications of this model application are clear, we recognize that trait-based datasets are inherently constrained by uneven natural history information across taxa, and some variables relevant to invasion (e.g., microbiome interactions, behavioral plasticity) remain difficult to quantify at scale. Additionally, factors such as genetic diversity are likely important to invasibility but are difficult and expensive to quantify and fall beyond the scope of this project. We also acknowledge that each trait was defined qualitatively and assessed through a literature review by three separate researchers. We addressed concerns about researcher bias by coordinating in-person meetings to clarify ambiguity and ensure continuity, and by having a single researcher review the entire matrix of species and trait values to maximize consistency of trait value definitions.

Similarly, using the number of invasive regions as a proxy for invasibility assumes comparable detection and reporting effort across regions, potentially underestimating invasion success for poorly monitored taxa. In turn, our dataset reflects the current literature for each species; thus, species that have traditionally been underrepresented in research are likely underrepresented in our dataset. Our model identifies variables that can predict arthropod invasibility, but it cannot address causality or define the relationship between the number of invaded regions and the predictor variables. Likewise, alternative frameworks (such as a Bayesian approach) may be selected in the future to understand these relationships causally or incorporate underlying phylogenetic matrices. While our model incorporates major life-history and environmental predictors, invasion outcomes are also shaped by trade networks, climate change, and stochastic events that are beyond the scope of trait-only analysis. In addition, we selected a single phylum to display invasive potential, but this application and methodology could be applied to other phyla with an appropriate dataset. The strong trait signals recovered here suggest that even with these limitations, biological characteristics alone contain meaningful predictive power. It should be noted that the true invasive potential of individual regions and species is highly context dependent. This model may be strengthened by integrating complementary approaches, such as species distribution models (SDMs), to predict regional suitability for potential invasion. However, SDMs may be poorly suited for taxa such as *Tropilaelaps* mites, as *Apis mellifera* can create buffered microhabitats that decouple mite survival from regional climate, suggesting that *T. mercedesae* may persist wherever its host occurs (Chantawannaku et al. 2018).

This methodology provides a basis for future work on traits and aspects important for predicting highly invasive species. This tool is invaluable for resource management and prioritization of limited assets involving invasive species and potential invaders. To ensure broad accessibility and transparency, the dataset curated for this work is publicly available on Harvard Dataverse and in Supplementary File 3. Future work can expand on this framework by integrating trait-based models with pathway-specific introduction pressure, climate-matching projections, and network models of global trade. Such multimodal approaches would allow forecasting not only *which* taxa are likely to become invasive, but also *where*, *when*, and *by which* routes they are most likely to spread. As global connectivity increases, developing anticipatory tools of this kind will be crucial for improving early-warning systems across agricultural, ecological, and public health contexts. Additionally, genomic predictors such as genetic diversity, gene count, and gene plasticity, which were largely unavailable at the time, may be important for predicting the invasive potential of arthropods (Stepien et al. 2005; Lawson Handley et al. 2011; Wellband et al. 2017).

In conclusion, our comparative model highlights generalizable traits associated with arthropod invasion success. Furthermore, our results identify *T. mercedesae* as a high-risk species that has already invaded many countries and has substantial potential to establish more invasive populations if introduced outside its current range. By leveraging existing data to predict future threats, trait-based models can help shift invasive species management from a reactive to a preventative paradigm — improving our ability to protect ecosystems, safeguard agriculture, and reduce economic impacts before invasions occur. Collectively, our results indicate a serious emerging threat to global apiculture and economies, calling for immediate investment in proactive defenses and research on *Tropilaelaps* mites.

## Statements and Declarations

## Funding

The authors declare that no funds, grants, or other support were received during the preparation of this manuscript.

### Authorship Credit

All authors contributed to the study conception and design. Material preparation and data collection were performed by Carmen Black, Treson Thompson and Madison Sankovitz. Statistical Analysis was performed by Carmen Black. Visuals were created by Madison Sankovitz and Carmen Black. The first draft of the manuscript was written by all authors and all authors commented on previous versions of the manuscript. All authors read and approved the final manuscript. Mentorship was provided by Samuel Ramsey.

## Supporting information

Supplementary Material One (Figure 1 References and Model Checks)

Supplementary Material Two (Parameter Defining)

## Acknowledgements

We would like to extend our gratitude to the University of Colorado Boulder and the Ramsey Research Foundation for supporting our research and hosting the Boulder Bee Lab.

Furthermore, we are grateful for the dedication of researchers and organizations that have dedicated their time and expertise to creating accessible data, including NOAA, GISD, CABI, GBIF, and many individuals who allowed us to create our own dataset. We would also like to thank Miles Moore at the University of Colorado and Chris Brundige for their support in the statistical elements of this paper, and Lincoln Taylor and Benjamin Young for reviewing our manuscript. Lastly, our team would like to recognize and thank all our international collaborators who enabled the collection of data on *Tropilaelaps* mites, including Aubrey Padilla, Mowgliz Productions, Center for Sustainability PH, and Gary Ayuste.

