## Supplementary Material One (Figure 1 References and Model Checks) for "A trait-based framework for quantifying arthropod invasion potential: Predictive modeling with *Tropilaelaps* mites as a case study"

Journal: *Biological Invasions*

Authors: Carmen Black, Treson Thompson, Madison Sankovitz, and Samuel Ramsey

Affiliation of the corresponding author: Department of Ecology and Evolutionary Biology, University of Colorado Boulder

### Figure 1 references

**Afghanistan:** Chantawannakul et al. 2016; de Guzman et al. 2017

**Azerbaijan:** Franco et al. 2025

**Bangladesh:** personal observation, 2024

**Cambodia:** Guerin et al. 2025

**China:** Anderson & Morgan 2007; Chantawannakul et al. 2016; Mohamadzade Namin et al. 2024

**Georgia:** Janashia et al. 2024

**India:** Anderson & Morgan 2007; Chantawannakul et al. 2016; Mohamadzade Namin et al. 2024

**Indonesia:** Anderson & Morgan 2007; Chantawannakul et al. 2016; Mohamadzade Namin et al. 2024

**Iran:** Mohamadzade Namin et al. 2024

**Kazakhstan:** Халыкова et al. 2025

**Laos:** Chantawannakul et al. 2016; Mohamadzade Namin et al. 2024

**Malaysia:** Anderson & Morgan 2007; Chantawannakul et al. 2016; Mohamadzade Namin et al. 2024

**Myanmar:** Chantawannakul et al. 2016; MinOo et al. 2023

**Nepal:** Chantawannakul et al. 2016; Mohamadzade Namin et al. 2024

North Korea

**Pakistan:** Chantawannakul et al. 2016; Mohamadzade Namin et al. 2024

**Papua New Guinea:** de Guzman et al. 2017; Mohamadzade Namin et al. 2024

**Philippines:** Anderson & Morgan 2007; Chantawannakul et al. 2016; Mohamadzade Namin et al. 2024

**Russia:** Joharchi & Stolbova 2024; Brandorf et al. 2025

**Singapore:** personal observation, 2023

**South Korea:** Chantawannakul et al. 2016; de Guzman et al. 2017; Mohamadzade Namin et al. 2024

**Sri Lanka:** Anderson & Morgan 2007; Mohamadzade Namin et al. 2024

**Tajikistan:** Franco et al. 2025

**Thailand:** Anderson & Morgan 2007; Chantawannakul et al. 2016; Mohamadzade Namin et al. 2024

**Ukraine:** GOV.UK 2025; Scheerlinck 2025

**Uzbekistan:** Mohamadzade Namin et al. 2024

**Vietnam:** Anderson & Morgan 2007; Chantawannakul et al. 2016; Mohamadzade Namin et al. 2024

Anderson, D. L., & Morgan, M. J. (2007). Genetic and morphological variation of bee-parasitic *Tropilaelaps* mites (Acari: Laelapidae): new and re-defined species. Experimental and Applied Acarology, 43(1), 1-24.

Brandorf, A., Ivoilova, M. M., Yañez, O., Neumann, P., & Soroker, V. (2025). First report of established mite populations, *Tropilaelaps mercedesae*, in Europe. Journal of Apicultural research, 64(3), 842-844.

Chantawannakul, P., de Guzman, L. I., Li, J., & Williams, G. R. (2016). Parasites, pathogens, and pests of honeybees in Asia. Apidologie, 47(3), 301-324.

de Guzman, L. I., Williams, G. R., Khongphinitbunjong, K., & Chantawannakul, P. (2017). Ecology, life history, and management of *Tropilaelaps* mites. Journal of economic entomology, 110(2), 319-332.

Janashia, I., Uzunov, A., Chen, C., Costa, C., & Cilia, G. (2024). First report on *Tropilaelaps mercedesae* presence in Georgia: The mite is heading westward!. Journal of Apicultural Science, 68(2), 183-188.

Joharchi, O., & Stolbova, V. V. (2024). The first report on the ectoparasitic genus *Tropilaelaps* Delfinado & Baker (Acari: Mesostigmata: Laelapidae) in Russia. Persian Journal of Acarology, 13(3), 513-516.

Franco, S., Laurent, M., & Véronique, D. (2025). Geographical Spread of the Exotic Mite *Tropilaelaps* spp.: State of Play of the Worldwide Situation in March 2025. Scientific Note published on the website of the European Union Reference Laboratory for Bee Health.

Guerin, E., Chheang, C., Sinpoo, C., Attasopa, K., Noirungsee, N., Zheng, H., ... & Disayathanoowat, T. (2025). Current Status, Challenges, and Perspectives in the Conservation of Native Honeybees and Beekeeping in Cambodia. Insects, 16(1), 39.

GOV.UK, 6 October 2025: Tropilaelaps mite restrictions: Ukraine.

MinOo, H., Kim, D. W., Akongte, P. N., Oh, D. G., Son, M. W., Park, B. S., ... & Choi, Y. S. (2023). Morphological Prevalence on Characterization of Varroa Mites from Myanmar. Journal of Apiculture, 38(4), 345-351.

Mohamadzade Namin, S., Joharchi, O., Aryal, S., Thapa, R., Kwon, S. H., Kakhramanov, B. A., & Jung, C. (2024). Exploring genetic variation and phylogenetic patterns of *Tropilaelaps mercedesae* (Mesostigmata: Laelapidae) populations in Asia. Frontiers in Ecology and Evolution, 12, 1275995.

Scheerlinck, J. P. (2025). Deadlier than varroa, a new honey-bee parasite is spreading around the world. The Conversation.

Халыкова, Г., Молдахметова, Г. А., Нұралиева, У. А., Крупский, О. Б., & Шералиева, Ж. Е. (2025). *APIS MELLIFERA* L КОЛОНИЯЛАРЫНДА *TROPILAELAPS* SPP МЕН *VARROA DESTRUCTOR*-ДЫҢ БІР УАҚЫТТА ТАРАЛУЫ ЖӘНЕ КӨБЕЮІ. Наука и образование, 5(2 (79)), 153-160.

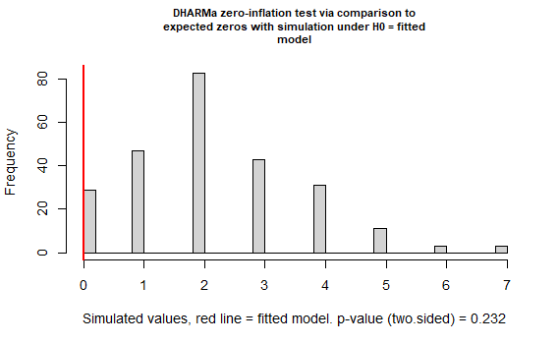

**Figure S1** DHARMa Zero Inflated test of the top performing model. No significant zero inflation detected in the model. Created in RStudio via DHARMa package.

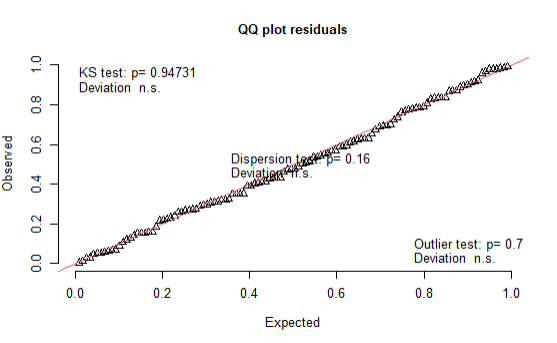

**Figure S2** DHARMa QQ Plot test of the top performing model. No significant outliers, dispersion, or deviation detected in the model.Created in RStudio via DHARMa package.

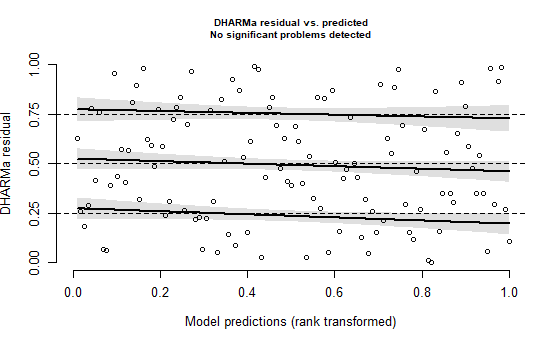

**Figure S3** DHARMa test results for residuals vs. fitted covariance of the top performing model. No significant problems detected for collinearity. Created in RStudio via DHARMa package.
