## Supplementary Material Two (Parameter Defining) for "A trait-based framework for quantifying arthropod invasion potential: Predictive modeling with *Tropilaelaps* mites as a case study"

Journal: *Biological Invasions*

Authors: Carmen Black, Treson Thompson, Madison Sankovitz, and Samuel Ramsey

Affiliation of the corresponding author: Department of Ecology and Evolutionary Biology, University of Colorado Boulder

**Native Continent(s)** (native_continent)

- The organism’s continent of origin. One of: Europe, Asia, Oceana, the Americas, Australia, Africa.
- Can be multiple

**Abundant in Native Range** (abundant_in_native_range)

- **Yes**: the organism is common in native regions, has high relative abundance, or is found in over 50% of the region. May be considered a pest or nuisance.
- **No**: the organism is uncommon in native range, has relative low abundance, or is found in less than 50% of the region.

**Past Intentional Introduction** (intentional_introduction)

- **Yes**: this organism was purposely introduced into its invasive range at least once in recorded history.
- **No**: the spread of this organism was never purposeful and happened by accident.

**Human Association** (human_association)

- **Very High:** the organism has a cosmopolitan distribution or is an important crop, medicine, companion animal, agricultural component, pollinator, etc., or is a parasite of any of the former. This may be an organism whose spread is dependent on humans. Obligate human parasites fall into this category.
- **High:** the organism is often highly sought after for its use as a traditional medicine, as an ornamental crop, for aesthetic values, cultural practices, or it is associated/obligate with organisms of the former. May be symbiotic with humans but humans are not an obligate host. Spread has likely been aided by human movement and trade.
- **Moderate:** the species is accidentally moved by humans, is sought after for various reasons in small geographic regions, provides some anthropogenic benefits, or is an obligate symbiont of the former. It may be aided by human trade but has other successful means of dispersal.
- **Low:** the organism is only occasionally moved by humans, is rarely sought by humans, or if it is, only in a small geographic context. Not a human symbiont or symbiont of a highly traded, anthropologically important species.
- **Very Low:** the species never sought directly by humans and has no known human benefits or connections. Dispersal is not facilitated by humans.

**Dormant Phase** (has_dormant_phase)

- **Yes:** the species has a life stage that is dormant and can survive otherwise harsh conditions through hibernation, suspended metabolism, pupation, incubation, or otherwise pausing normal activity in order to survive until a more favorable environment is encountered.
- **No:** all life stages must remain metabolically active and cannot survive long periods of food deprivation or extreme temperature or moisture.

**Disease Vector** (vector)

- **Yes:** the species is capable of transmitting pathogens or parasites mechanically or biologically in any capacity. This includes both human and non-human pathogens that may affect the organism’s host or other associated organisms.
- **No:** the species does not vector any known pathogens that affect humans, its host, or associated organisms. Species with this designation can transmit diseases between conspecifics as long as the pathogen only affects this species.

**Dispersal Capacity** (dispersal_capacity)

- **Very High:** this species has spread rapidly over a short period of time or has a cosmopolitan distribution. Has a life stage adapted for long range dispersal, a host with a broad range, or otherwise possesses an extremely effective means of dispersal. For example, may be able to fly for long distances, exploit global ocean and wind currents, have a high affinity for human transportation infrastructure (such as water ballast of a cargo ship), be associated with a species with a broad range, or parasitize a species that is transported globally.
- **High:** this species has spread rather quickly and is found almost worldwide. Has a dispersal stage of life, a broadly dispersed host, or a means of dispersal that it frequently utilizes. May have another means of dispersal such as frequently being transported accidentally.
- **Moderate:** the organism is found in around ½ of the countries worldwide. May have a dispersal stage of life, mechanism, or host but is not prolific.
- **Low:** the organism is found in less than ½ countries worldwide and it is not aided by other species or human infrastructure. If it is dispersed in some way, its range is limited.
- **Very Low:** it is found in less than ½ countries worldwide. It does not have a dispersal stage of life, mechanism, or host. Its dispersal is not aided by other species or human infrastructure and its dispersal range is limited.

**Number of Native Regions** (number_native_regions)

- Count of the number of regions listed as native in literature, CABI database, and certified records. Only regions with a verified, established, historical population are included in this count.
- Regions are used instead of the political subdivisions of countries in order to equitably compare smaller and larger countries with equal numbers of political subdivisions. For example, provinces of large countries like Canada are counted as individual regions, while smaller countries like Vietnam are considered a single region.
- Regions are counted as native based on literature consensus about the ecological and historical context of the species.

Subdivided Countries: countries where each political subdivision (at the first hierarchical level; usually state, province, or department) is counted as an individual region.

| **Country** | **Justification for Subdivision** |
| --- | --- |
| Australia | Large size of provinces which contain different biomes. |
| Brazil | Large size of country which contains a vast distribution of biomes and climates. Distinct borders and controls for each province. |
| Canada | Large size of provinces. Distinct borders and controls for each province. |
| China | Large size of country which contains a vast distribution of biomes and climates. Distinct borders and controls for each province. |
| India | Large size of country which contains a vast distribution of biomes and climates.Distinct borders and controls for each state and union territory. |
| Indonesia | The country is composed of many islands, many of which are ecologically isolated from each other. |
| Japan | The country is composed of many islands, many of which are ecologically isolated from each other. |
| Malaysia | The country is composed of many islands, many of which are ecologically isolated from each other. |
| Philippines | The country is composed of many islands, many of which are ecologically isolated from each other. |
| Russia | Large size of country and different biomes contained in each region. |
| United States | Large size of states which contain drastically different biomes. Distinct borders and controls for each state. |

Partially Subdivided Countries: countries that are considered a single region except for the listed island which is isolated from the mainland and ecologically distinct. Examples:

| **Country** | **Separate Region(s)** |
| --- | --- |
| Spain | Canary Islands |
| Portugal | Madeira |
| Ecuador | Galápagos Islands |

Single-region Countries: countries not previously listed are considered a single region.

**Number of Introduced Regions** (number_introduced_regions)

- Count of the number of regions listed as introduced in primary literature, CABI database, and certified records. Only regions with verified reports of established populations are included in this count, but quantification of the ecological impact of these populations is not required for inclusion. Does not include incidental reports that have not been verified or populations that have been eradicated.
- See above tables for definitions of a region.

**Inconspicuousness** (easy_to_overlook)

- **Yes**: organisms that are small, adept at hiding, or easily mistaken for another organism, or have a brief life stage matching this description (i.e. adults are well camouflaged but eggs are bright, conspicuous, and hatch quickly). A layperson would not be able to identify this species or life stage and an expert would have difficulty identifying it.
- **Sometimes:** organisms that have only one small and easy to overlook part of its life cycle (i.e. nondescript eggs that hatch quickly). This organism could be overlooked by a layperson but is easily spotted by an expert.
- **No**: all stages are conspicuous and a layperson would notice this species, although may not be able to identify it. An expert would immediately recognize its presence and identity.

**Physiological Tolerance** (physiologic_tolerance)

- **Very high:** the organism is able to survive all or almost all climate types, often being able to live through freezing, drought, and flooding conditions. Shows extremely high tolerance to environmental conditions and likely found in most climate types across the globe. Includes euryhaline organisms. May have unique adaptations that allow it to survive diverse environmental conditions.
- **High:** this category falls between very high and moderate. Includes organisms that are found in at least three major climate types and two subtypes. These organisms may have a wider range of tolerance to temperature extremes, precipitation, humidity etc than moderate. This may be an organism that tolerates brackish water and freshwater or saltwater.
- **Moderate:** this category includes organisms that are found in at least one other climate range that is not a sub-climate of their native climate. These organisms often are intolerant to temperature extremes and are limited by annual precipitation to some extent. If aquatic, they are not euryhaline.
- **Low:** organisms that are found in their native climate range and a sub-climate of their native climate (ex tropical and subtropical). They have distinct temperature ranges and precipitation requirements for life. Can tolerate relatively small changes in temperature, salinity, altitude, etc.
- **Very Low:** organisms that are only able to tolerate one climate type and have strict environmental conditions to sustain life. Cannot tolerate change in their environment

**Reproductive Capacity** (reproductive_capacity)

- **Very high,** organisms that have continuous reproduction once reproductive age is achieved and environmental conditions allow. These are R selected species that spend most of their adult life mostly reproducing, often having large numbers/ multiple offspring at once. This may also include organisms that can reproduce via parthenogenesis (without mating or sexual reproduction). This may also include species that can have large clutches of offspring and produce multiple clutches per reproductive season. This almost always includes eusocial organisms that have or can have continual reproduction via a single or multiple queens. Other reproductive factors may influence ranking.
- **High,** organisms that reproduce multiple times per year or reproductive season for all of their reproductive age. Often R selected species that have large numbers/ multiple offspring at once. This likely includes species that are able to reproduce multiple clutches in a single reproductive season and/or likely to reproduce a large number of offspring in a single reproductive season. Other reproductive factors may influence ranking.
- **Moderate,** organisms that produce roughly once a year (annually) or once per reproductive season once reaching reproductive age. They may have a large number of offspring at once but only have one mating/reproductive event in a single year. These organisms can not reproduce via parthenogenesis.
- **Low,** organisms that reproduce less than once a year or reproductive season once establishing reproductive age (reproducing every 2,3,4 years). These organisms can not reproduce via parthenogenesis. These organisms could possibly reproduce once a year, but if they do they produce < 100 offspring per year.
- **Very Low,** organisms that reproduce roughly less than every 5 years or 5 reproductive seasons once establishing reproductive age. (reproducing every 5+ years). This strategy is very uncommon for arthropods. They may reproduce less than once a year or reproductive season once establishing reproductive age (reproducing every 2,3,4 years) but reproduce low numbers of offspring (< 100).

**Maximum Clutch Number per Year** (max_number_clutch_per_female_annual)

- Use primary literature to find how many clutches a single reproductive female can reproduce. Use the highest peer-reviewed documentation available. A clutch may be single depending on the species (i.e. chrysoperla lay single eggs not clusters); if so this should be the maximum recorded number of eggs produced in the year (i.e. highest annual fecundity).

**Maximum Offspring per Clutch** (max_number_offspring_per_clutch)

- Use primary literature to find how many eggs a single reproductive female can reproduce in a clutch. Use the highest peer-reviewed documentation available. A clutch may be single depending on the species (i.e. chrysoperla lay single eggs not clusters); if so this should be one. This does not include relative survivorship.

**Maximum Generations per Year** (max_number_ generation_per_year)

- Use primary literature to find how many complete generations can occur in a single year. Use the highest peer-reviewed documentation available in their native or introduced range. If a species can complete half a generation round down to the nearest number for complete generation (i.e. to adult). For example, in ideal climate an arthropod can produce 6.5 generations; this would be recorded as 6 complete generations.

**Small Population Resilience** (small_population_resilance)

- **Very high:** organisms that have strong genetic diversity and effective gene flow, robust ecological or behavioral adaptations to disturbance, are parthenogenic, or otherwise able to avoid local extirpation when population size is only a few individuals.
- **High:** organisms that have moderate to high genetic diversity, may have some natural or human-aided population connectivity, can recover from moderate disturbances, or are otherwise able to avoid local extirpation when population size is less than one hundred.
- **Moderate:** organisms that have moderate genetic diversity, have limited connectivity to other populations, are vulnerable to more intense or frequent disturbances, or will locally extirpate if the population is less than one hundred individuals.
- **Low:** organisms with low genetic diversity (inbreeding), are vulnerable to disturbance, have isolated populations, and populations extirpate when there are few individuals.
- **Very Low:** organisms that have critically low genetic diversity, have extremely isolated populations, and populations would require significant intervention in order to avoid extirpation or extinction.

**Ontogeny** (ontogeny)

- **Simple:** animals that develop through a sequential and remain morphologically constant. The juvenile stage closely resembles the adult, with the main difference being size and sexual maturity. Includes mammals and hemimetabolous insects, for example.
- **Complex:** animals that develop through developmental stages that are morphologically distinct, such as a larval or pupal stage. Juveniles and adults occupy different ecological niches. Includes most amphibians, holometabolous insects, and many marine invertebrates.

**Diet breadth-** The following categories were answered yes or no for each organism (only one of the three categories can be selected)

- **Monophagic:** organisms that only have one food source or require a specific host to complete its life cycle. (Yes or No)
- **Oligophagic:** organisms that can feed on multiple species of the same category. (Yes or No)
- **Polyphagic:** organisms that can derive their nutritional needs from a broad spectrum of species and sources. (Yes or No)

**Lower Mean Annual Temperature Threshold** (mean_annual _temp_lower_C)

- Literature consensus on the lowest temperature threshold that the organism can survive, including adaptations for niche construction (i.e. a wood boring beetle that can survive extremely cold winters by constructing galleries within trees).
- If no research has been conducted on thermal thresholds, the lowest mean temperatures for the organism’s native and introduced regions was calculated using the National Weather Service’s Climate Prediction Center.
  - To calculate the mean lower temperature known regions were surveyed and annual low temperatures were compiled and averaged from all documented ares with a weather station.

**Higher Mean Annual Temperature Threshold** (mean_annual _temp_upper_C)

- Literature consensus on the highest temperature threshold that the organism can survive, including adaptations for niche construction (i.e. a termite colony that constructs ventilation systems).
- If no research has been conducted on thermal thresholds, the highest mean temperatures for the organism’s native and introduced regions was calculated using the National Weather Service’s Climate Prediction Center.
  - To calculate the mean upper temperature known regions were surveyed and annual high temperatures were compiled and averaged from all documented areas with a weather station.

**Native Climate** (native_climate)

- Choose from major tropical, subtropical, temperate, continental, dry, and polar
  - Or subtypes if the organism is more niche that may include mediterranean, semi-arid, arid, subtropical, subarctic, desert, humid subtropical, tundra, etc.
  - Subtypes can be variable and categorization of climate types is variable and will not be included in analysis.

**Introduced/Invasive Climate** (invasive_climate)

- Choose from major tropical, subtropical, temperate, continental, dry, and polar
  - Or subtypes if the organism is more niche that may include mediterranean, semi-arid, arid, subtropical, subarctic, desert, humid subtropical, tundra, etc.
  - Choose all that apply to invasive range

**Average Higher Annual Precipitation in mm** (native_avg_precip_upper_(mm))

- Use primary literature to find precipitation thresholds; if not researched use:
- For the Categories Below please use Climate Prediction Center (National Weather Service) <https://www.cpc.ncep.noaa.gov/products/timeseries/> Here you can select temperature or precipitation then zoom in to the weather station from the reported native region. Here you can click on the 365 days for both temperature and precipitation then can calculate the mean temperature and precipitation over the last 12 months (one temperature or precipitation per each month).

**Average Lower Annual Precipitation in mm** (native_avg_precip_lower_(mm))

- Use primary literature to find precipitation thresholds; if not researched use:
- For the Categories Below please use Climate Prediction Center (National Weather Service) <https://www.cpc.ncep.noaa.gov/products/timeseries/> Here you can select temperature or precipitation then zoom in to the weather station from the reported native region. Here you can click on the 365 days for both temperature and precipitation then can calculate the mean temperature and precipitation over the last 12 months (one temperature or precipitation per each month).

**Natural Predators in Native Range** (native_has_natural _predators)

- Are there natural predators in native range? If so do native predators help control the population?
- Are these somewhat limiting to the organism’s success?
- Yes or no- If no for any then no for column

**Natural Predators in Introduced/Invasive Range (**Invasive_has_natural _predators)

- Are their natural predators in the introduced range? Or is it able to outcompete native species because of a lack in predation.
- Are these somewhat limiting to the organism’s establishment in the introduced range?
- Yes or no- If no for any then no for column

**Upper Altitude Limit** (altitude_upper_(m))

- Use primary literature to find precipitation thresholds; if not researched use. The highest altitude where the organism is found. For aquatic animals, record water depth regardless of absolute altitude.

**Lower Altitude Limit** (altitude_lower_(m))

- Use primary literature to find altitude thresholds. The lowest altitude where the organism is found. For aquatic animals, record water depth regardless of absolute altitude, i.e. for aquatic animals this value should be zero.

**References and Citations for Biological and Ecological Traits**  (new_sources) (Reference)

- All citations here for all literature used to collect data.
